# Subtractive Genomic Analysis for Identification of Novel Drug Targets and Vaccine Candidates against *Bartonella bacilliformis* subsp. Ver097

**DOI:** 10.1101/731570

**Authors:** Md. Tahsin Khan, Araf Mahmud, Md. Asif Iqbal, Mahmudul Hasan

## Abstract

*Bartonella bacilliformis* is the causative agent of Carrión’s disease, one of the truly neglected tropical diseases found in Peru, Colombia and Ecuador. Recent evidence predicts that *Bartonella bacilliformis subsp*. ver097 can emerge as an antibacterial resistant strain and hence identification of novel drug targets is a crying need. Subtractive genome analysis of *B. bacilliformis* subsp. ver097 was successfully done in order to address the challenges. Various computational tools and online based servers were used to screen out human homologous proteins of pathogen and proteins involved in common metabolic pathways of host and pathogen. Only 7 proteins involved in pathogen specific pathways were further analyzed to identify membrane proteins. ‘Flagellar biosynthesis protein FlhA’ and ‘ABC transporter permease’ were found to be novel as targets according to DrugBank database. To avoid side effects in human while administering drugs, human ‘anti-targets’ analysis was performed to confirm non-homology of selected novel drug targets. Both predicted proteins also showed probability of antigenicity prediction through VaxiJen, however, ‘Flagellar biosynthesis protein FlhA’ showed broad spectrum conservancy with *Bartonella* strains. Therefore, Flagellar biosynthesis protein FlhAcould facilitate the development of novel drugs and therapeutic compounds along with vaccines for efficient treatment of infections caused by *Bartonella bacilliformis subsp.* ver097.

## Introduction

*Bartonella bacilliformis*, a gram-negative, intracellular, facultative, aerobic[1] and a member of the alpha-2 subgroup Proteobacterium is the causative agent of Carrión’s disease in humans. This pathogen is highly motile by 3 to 10 micrometers long polar lophotrichous flagella and appendages consisting of approximately 42kDa flagellin protein subunits[2]. Carrión’s disease has been reported as an endemic in the river valleys of Peru, Ecuador as well as Colombia[3, 4]. However, Peru is considered as the most important endemic area for Bartonellosis[5]. The only known natural reservoir for this pathogen is human. *Bartonella bacilliformis* is believed to be transmitted by the bites of a sand fly, *Lutzomyiaverrucarum* as well as other possible closely related sand flies[4].

Carrion’s disease can occur in two clinical forms; verruga peruana-generally is a non-fatal form characterized by nodular cutaneous eruptions, and the other is Oroya fever, a frequently fatal disease in absence or delay of antibiotic treatments[2]. Verruga-peruana generally persists 3 to 6 months and wart-like hemangiomatous lesions are produced in the skin along with mucous membranes due to the entrance of the bacteria in the endothelial cells. The lesions are classified as miliary (small reddish papules less than 3 mm in diameter), mular (nodular tumors greater than 5 mm in diameter) and diffuse subdermal nodules[6]. However, Oroya fever, commonly found in children [8] is a life-threatening acute phase in which erythrocytes are destroyed by the spleen due to the entrance of bacteria in erythrocytes[7], and it has a high fatality rate (up to 88%) while untreated [8]. General malaise, myalgia, headache, hepato/splenomegaly, palor, jaundice besides fever are the main symptoms of this phase[3]. Some patients may develop a transient immunosuppression, as a consequence of the septic state which results in the eve of other opportunistic infections, for example, toxoplasmosis, tuberculosis, salmonellosis, malaria, shigellosis, histoplasmosis and, pneumoctytocis phase [3, 9, 10].

The majority of pathogens causing infections and diseases are resistant to more than one drug[11]. Though studies suggested that most of *Bartonella* species including *B. bacilliformis* are highly susceptible to mostof antibiotics, such as beta-lactams, aminoglycosides, chloramphenicol, rifampin, macrolides, tetracyclines, cotrimoxazole, fluoroquinolones[12] and gentamicin[1], but there is a potential risk of the spontaneous selection of successful antibiotic resistant *B. bacilliformis* clone in a clinical setting. For examples, few studies developed in vitro has showed the ability of *B. bacilliformis* to acquire chromosomal mutations leading to high levels of antibiotic resistance [13,14]. Lack of comprehensive analysis of *B. bacilliformis* genomic structure is an obstacle in developing species-specific new therapeutic compounds and vaccine candidates [5].

In recent years, identification of species-specific drug targets and vaccine candidates has been performed on several pathogenic strains through subtractive genomic approaches [15-18]. Analysis of mutational divergence of core genes revealed the emergence of multiple sub-species or outlier for *B. bacilliformis* [19, 20]. In the study, proteome of *Bartonella bacilliformis* supsp. ver097 was retrieved for applying subtractive genomic approaches. Various computational tools were used to identify proteins that are essential for the survival of the pathogen. Host non-homology analysis, metabolic pathways analysis were performed to avoid both cross-reactivity of drugs with human proteins and involvement of pathogenic proteins in host metabolism respectively. This study was further augmented to identify membrane proteins showing novelty as drug targets. Antigenicity testing was performed to check whether the identified novel drug targets. The study would be helpful to develop effective broad spectrum drugs target and vaccine candidates against *Bartonella bacilliformis.*

## Materials and methods

The whole proteome *of Bartonella bacilliformis subsp.* ver097was analyzed according to subtractive genomic approach to recognize novel drug targets as well as vaccine candidates. The overall workflow is illustrated in Figure 1.

**Figure 1:**
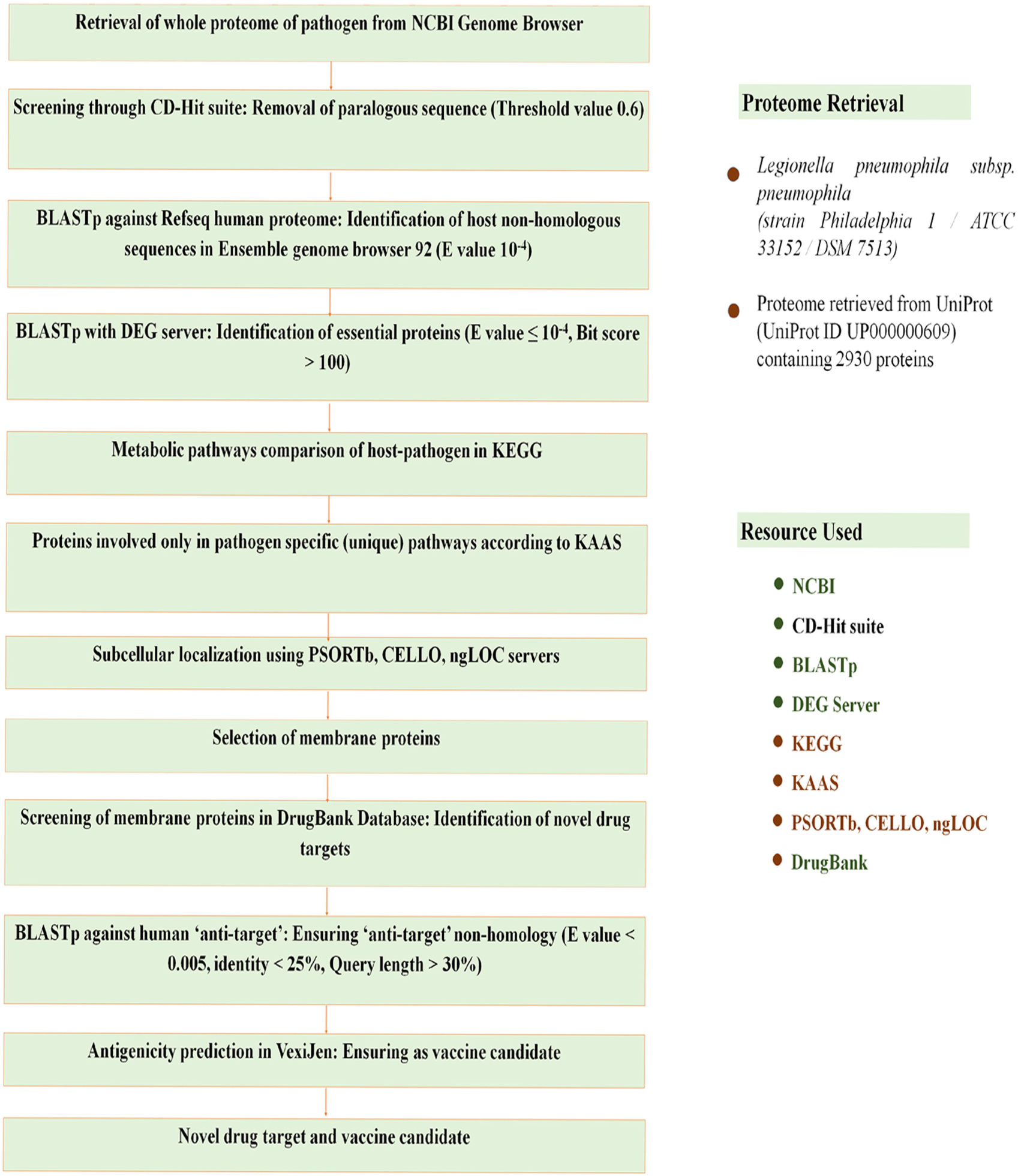
An outline of Subtractive Genomics approach to Bartonellabacilliformis subsp.Ver097 proteomes for identification of novel drug candidates.

### Retrieval of Complete Proteome (Set0)

The whole proteome of *B. bacilliformis* subsp. ver097 (Assembly GCF_000709795.1) was retrieved from NCBI Genome database (https://www.ncbi.nlm.nih.gov/genome).

### Identification of Paralogous Sequences (Set1)

To remove the paralogous sequences from the *B. bacilliformis* subsp. ver097 proteome, the protein sequences of set0 were subjected to CD-Hit suite[21](http://weizhongli-lab.org/cdhit_suite/cgi-bin/index.cgi?cmd=cd-hit). The server is programmed to take input as a fasta format sequence database which produces output as a set of non-redundant as well as representative sequences. To eliminate redundant sequences through this program, cutoff score 0.6 of sequence identity was set. Paralogous proteins with more than 60% identity were excluded from set0. Moreover, proteins <100 amino acids were excluded from set0. The remaining proteins were listed as set1 as non-paralogous protein sequences.

### Identification of Protein Sequences Non-homologous to the Proteome of Human (Set2)

The aim of this step was to avoid functional similarity with human proteome in order to prevent bindings of therapeutic compounds to the active site of the host homologous proteins. Non-paralogous proteins of the pathogen were analyzed using BLASTp against human Refseq proteome in Ensemble Genome Database (https://uswest.ensembl.org/Multi/Tools/Blast?db=corE)[22]. Proteins of the pathogen were considered host homologous proteins if any significant hit above the threshold value 10^−4^ was found. These homologous proteins were filtered while non-homologous proteins were listed as set2.

### Identification of Essential Non-homologous Proteins (Set3)

Host non-homologous proteins of the pathogen listed in set2 were subjected to the Database of Essential Genes (DEG)[23]. DEG15.2 server contains 53,885 essential genes along with 786 essential non-coding sequences. By selecting all the stains of organisms present DEG 15.2 server, BLASTp was carried out using threshold value 10^−10^ and minimum bit score 100 as parameters. Proteins hit with expectation value ≤10^−100^, identity ≥ 25% were listed in set3considering as essential proteins of the pathogen. Products of essential genes constitute a minimal genome for a bacterium, that play key roles in the field of synthetic biology[24].

### Analysis of Metabolic Pathway (Set4-a and Set-b)

Metabolic pathways of *B. bacilliformis* subsp. ver097were analyzed against the human metabolic pathways through KEGG server[25]. KEGG is Kyoto Encyclopedia of Genes and Genomes which contains complete metabolic pathways present in living organisms. All metabolic pathways of *B. bacilliformis* and host were collected from KEGG PATHWAY Database[26] using their respective three letter KEGG organism code; ‘bbk’ for *B. bacilliformis* and ‘has’ for *H. sapiens*. A comparison was made in order to recognize the unique metabolic pathways only present in the pathogen, while the remaining pathways of the pathogen were grouped as common.

Identified host non-homologous, essential proteins of *B. bacilliformis* subsp. ver097were subjected to BLASTp through KAAS server[27] at KEEG database. Proteins present only in the unique metabolic pathways of the pathogen were listed as set4-a and proteins involved in common pathways or only in host pathways were excluded. KAAS provides functional annotation of genes through a blast comparison against the manually curated database of KEGG GENES. KO (KEGG Orthology) assignments specify metabolic proteins. KAAS also generates KEGG pathways which indicate these metabolic proteins. Proteins having no KO assignment were listed as set4-b.

### Prediction of Subcellular Localization of set4-a

Since *B. bacilliformis* subsp. ver097is a gram-negative bacterium, proteins of this bacterium can be recognized from five feasible subcellular locations. These are cytoplasm, inner membrane, periplasm, outer membrane and extracellular space. Proteins functioning in cytoplasm can be used as putative drug targets while surface membrane proteins can be considered both as drug targets and vaccine candidates. To predict subcellular localization of shortlisted proteins of set4-a, PSORTb v3.0.2 (http://www.psort.org/psortb/index.html) [28], CELLO v.2.5[29] (http://cello.life.nctu.edu.tw/), ngLOC[30] (http://genome.unmc.edu/ngLOC/index.html) servers were used. The best score generated by these servers for a location was counted. Final location of the proteins was fixed by considering the reported location of more than two servers. Only membrane proteins were listed in set5 as potential drug targets and vaccine candidates.

### Identification Novel Drug Targets by Screening and Druggability Analysis (Set6-a and Set6-b)

Membrane proteins of set5 were further analyzed to check the novelty as drug target. Proteins were screened out through database of DrugBank 5.1.0 [31]while using default parameters. DrugBank 5.1.0 is a database containing entries including 2556 approved small molecule drugs, 5125 experimental drugs, 962 approved biotech drugs and 112 nutraceuticals. Moreover 511 non-redundant protein sequences such as drug target, enzyme, carrier or transporter are linked to these drug entries. Presence of drug targets in the database indicating the same function was listed as druggable targets as set6-a, while the absence of drug entries for a protein indicated its novelty as a drug target. Since our aim is to identify novel drug targets for *B. bacilliformis* subsp. ver097, proteins showing no similarity after screened through DrugBank database, were listed in set6-b.

### ‘Anti-target’ Analysis of Essential, Nonhomologous and Novel Drug Targets

Drugs or therapeutic compounds designed to bind and inhibit the action of a protein present in the pathogen may dock with some crucial proteins of host serving pharmacological effects in the host. These proteins of the host are termed as ‘anti-targets’. In human, ‘anti-targets’ include ether-a-go-go related gene (hGER), P-glycoproteins (P-gp), the pregene X receptor (PXR) and constitutive and rostane receptor (CAR). Additionally some membrane receptors, e.g. dopaminergic D2, adrenergicα_1a_, serotonergic 5-HT_2C_, and the muscarinic M1 are included as ‘anti-targets’. Total 210 ‘anti-targets’ were reported in the literature with their accession numbers[32]. The corresponding protein sequences were fetched from NCBI Protein database and these accession numbers are provided in Supplementary file SF1. Novel drug targets were subjected to BLASTp analysis in NCBI blast program against these human ‘anti-targets’ setting an E-value <0.005, query length >30%, identity <25% as parameters. Proteins showing <25% identity were listed as set7.

### Prediction of Antigenicity (Set8)

To predict the protective antigens as well as subunit vaccines, novel drug target proteins were screened through VaxiJen v2.0 server[33](http://www.ddg-pharmfac.net/vaxijen/VaxiJen/VaxiJen.html) using threshold value 0.4. Protein sequences showing antigenicity >0.4 were listed in set8. This server was developed in order to allow classification of antigen based on the physiochemical properties. VaxiJen is the first server to predict of protective antigens as alignment independent.

### Conservancy Analysis of Predicted Sequences with Other Strains

Conservancy pattern of the predicted sequences with other classically used strains is essential for determining the range of drug spectrum among the entire homologous bacterial community. Conservancy analysis of the predicted drug target sequences of was done by BLASTp suite of NCBI server (https://blast.ncbi.nlm.nih.gov/Blast.cgi?PAGE=Proteins). Here, in case of performing protein-protein BLAST, all of the parameters were kept default except *Bartonella bacilliformis* in organism option.

## Results and Discussion

Identification of novel drug targets, as well as vaccine candidates against *B. bacilliformis* subsp. ver097 was the prime concerned target of the study, and here, subtractive genomic analysis of the entire *B. bacilliformis* subsp. ver097 proteome was done by using different computational tools and database. A brief eyesight of the subtractive genomic approach was presented in Table 1.

**Table 1:** Subtractive genomic analysis scheme.

| *S. No.* | *Steps* | *B. bacilliformis* subsp. ver097 |
| --- | --- | --- |
| 1 | Total number of proteins | 1097 |
| 2 | Removed non-paralogous ( >60% identical) and smaller proteins (>100 amino acids) in CD-Hit and | 100 |
| 3 | Number of proteins nonhomologous to *H. sapiens* using BLASTp (E value $10^{-3}$) | 603 |
| 4 | Essential proteins in DEG 15.2 server (E value $\leq 10^{-100}$, Bit score >100) | 114 |
| 5 | Essential Proteins involved only in unique metabolic pathways (KAAS at KEGG) | 7 |
| 5 | Proteins assigned KO (KEGG Orthology) but not in any pathway | 27 |
| 6 | Essential membrane proteins using PSORTb, CELLO, ngLOC, | 4 |
| 7 | Essential proteins found to be novel in DrugBank 5.1.0 (using default parameters) | 2 |
| 8 | Novel drug target proteins non-homologous to ‘anti-targets’ using BLASTp (E value <0.005, Identity <25%, Query length >30%) | 2 |
| 9 | Proteins showing antigenicity using VaxiJen v2.0 (Threshold value >0.4) | 2 |
| 10 | Highly conserved protein | 1 |

### Identification of Paralogous Sequences

Entire proteome of *B. bacilliformis* subsp. ver097contains 1097 proteins. Paralogous protein sequences of the pathogen were identified through CD-hit server. Total 8 paralogous clusters were found among set0 proteins of the pathogen. The paralogous sequence above >60% similarity were removed leaving 1087 non-paralogous protein sequences. Furthermore, total proteins containing <100 amino acids were filtered out from 1087 non-paralogous protein sequences assuming these smaller proteins unlikely represent the essentiality[34,35]. Total 997 non-duplicate proteins which contain ≥100 amino acids in the sequences were listed in set1.

### Identification of Human non-homologous Proteins

Proteins found to be involved in common cellular systems emerged as homologous in course of time between bacteria and human[35-37]. As a result, drugs and therapeutic compounds developed to bind proteins of any pathogen should avoid unintentional blocking of host proteins. To avoid cross-reactivity with host proteins, all non-paralogous proteins of the pathogen (set1) were subjected to BLASTp against Refseq proteome of *Homo sapiens* in Ensemble genome browser 92. The result revealed that total 394 proteins were found to be homologous with the human proteins. These proteins were removed while 603 non-homologous proteins were listed in set2.

### Selection of Essential Proteins

The most promising drug targets are essential proteins because most antibacterial compounds are designed to dock in essential gene products[38]. Screening through DEG 15.2 server, 114 proteins were found to be similar above the parameters. These proteins were selected and listed in set3 as essential proteins for the survival of *B. bacilliformis* subsp. ver097. Proteins encoded by essential genes unique to an organism may be considered as species-specific therapeutic targets[39].

### Analysis of Metabolic pathway

KEGG server contains 95 metabolic pathways for *B. bacilliformis* subsp. ver097and 325 pathways for human. Through manual comparison, 24 metabolic pathways were found to be pathogen-specific and are provided in Supplementary Table 1. Proteins involved in these unique pathways can be selected to as drug targets.

Non-homologous essential proteins subjected to BLASTp in KAAS server at KEGG revealed that 82 proteins among 114 were assigned both KO (KEGG Orthology) and metabolic pathways. To avoid undesirable cross-reactions with human pathways, 75 proteins among 82 were excluded, leaving only 7 proteins involved in pathogen-specific pathways. These unique pathway proteins were listed in set4-a and are provided in Table 2. Only Set4-a proteins were used to identify novel drug targets and vaccine candidates in our study.

**Table 2:**
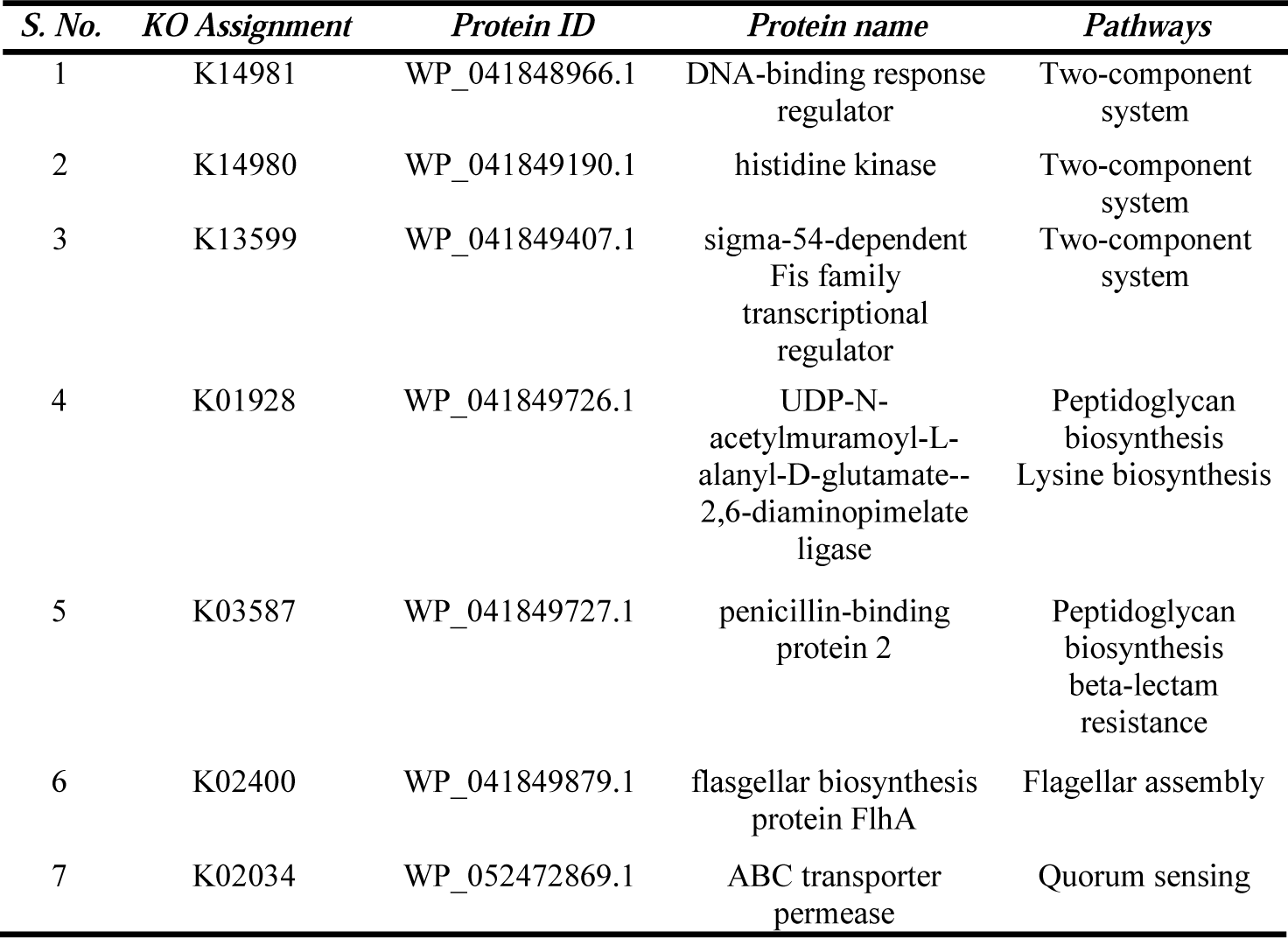
Metabolic proteins involved only in pathogen-specific pathways.

| *S. No.* | *KO Assignment* | *Protein ID* | *Protein name* | *Pathways* |
| --- | --- | --- | --- | --- |
| 1 | K14981 | WP_041848966.1 | DNA-binding response regulator | Two-component system |
| 2 | K14980 | WP_041849190.1 | histidine kinase | Two-component system |
| 3 | K13599 | WP_041849407.1 | sigma-54-dependent Fis family transcriptional regulator | Two-component system |
| 4 | K01928 | WP_041849726.1 | UDP-N-acetylmuramoyl-L-alanyl-D-glutamate--2,6-diaminopimelate ligase | Peptidoglycan biosynthesis<br>Lysine biosynthesis |
| 5 | K03587 | WP_041849727.1 | penicillin-binding protein 2 | Peptidoglycan biosynthesis<br>beta-lactam resistance |
| 6 | K02400 | WP_041849879.1 | flagellar biosynthesis protein FlhA | Flagellar assembly |
| 7 | K02034 | WP_052472869.1 | ABC transporter permease | Quorum sensing |

Among 114 proteins, 27 were found not to be involved in metabolic pathways but were assigned KO (KEGG Orthology). Since these proteins had successfully passed through host non-homology test and essentiality test, in future they can be used as drug targets as well as vaccine candidates. These 27 proteins are provided in Table 3. Only 5 proteins among 114 listed as set4-b were found not to be assigned KO meaning that they are not involved in any metabolic pathway of the pathogen and the host. They are provided in Table 4.

**Table 3:** Proteins assigned KO but not assigned any pathway.

| *KO assignment* | *Accession No.* | *Protein name* |
| --- | --- | --- |
| K03296 | WP_041848830.1 | AcrB/AcrD/AcrF family protein |
| K03466 | WP_041848841.1 | DNA translocase FtsK |
| K03089 | WP_041848844.1 | RNA polymerase sigma factor RpoH |
| K19689 | WP_041848873.1 | aminopeptidase |
| K02600 | WP_041848898.1 | transcription termination/ |
|  |  | antitermination protein NusA |
| K07391 | WP_041848977.1 | ATP-binding protein |
| K03496 | WP_041849016.1 | ParA family protein |
| K03497 | WP_041849017.1 | chromosome partitioning protein ParB |
| K03641 | WP_041849127.1 | translocation protein TolB |
| K03797 | WP_041849219.1 | S41 family peptidase |
| K03568 | WP_041849289.1 | metalloprotease TldD |
| K02483 | WP_041849309.1 | DNA-binding response regulator |
| K02198 | WP_041849313.1 | hemelyaseNrfEFG subunit NrfE |
| K04485 | WP_041849362.1 | DNA repair protein RadA |
| K07082 | WP_041849372.1 | endolytictransglycosylaseMltG |
| K03499 | WP_041849408.1 | Trk system potassium transporter TrkA |
| K07018 | WP_041849624.1 | alpha/beta hydrolase |
| K09014 | WP_041849626.1 | Fe-S cluster assembly protein SufB |
| K09456 | WP_041849647.1 | acyl-CoA dehydrogenase |
| K03761 | WP_041849702.1 | MFS transporter |
| K03086 | WP_041849735.1 | RNA polymerase sigma factor RpoD |
| K03498 | WP_041849746.1 | metal ABC transporter ATPase |
| K03924 | WP_041849766.1 | MoxR family ATPase |
| K09015 | WP_052472887.1 | Fe-S cluster assembly protein SufD |
| K03820 | WP_080772740.1 | apolipoprotein N-acyltransferase |
| K04744 | WP_080772760.1 | LPS-assembly protein LptD |
| K00528 | WP_041849825.1 | ferredoxin--NADP reductase |

**Table 4:**
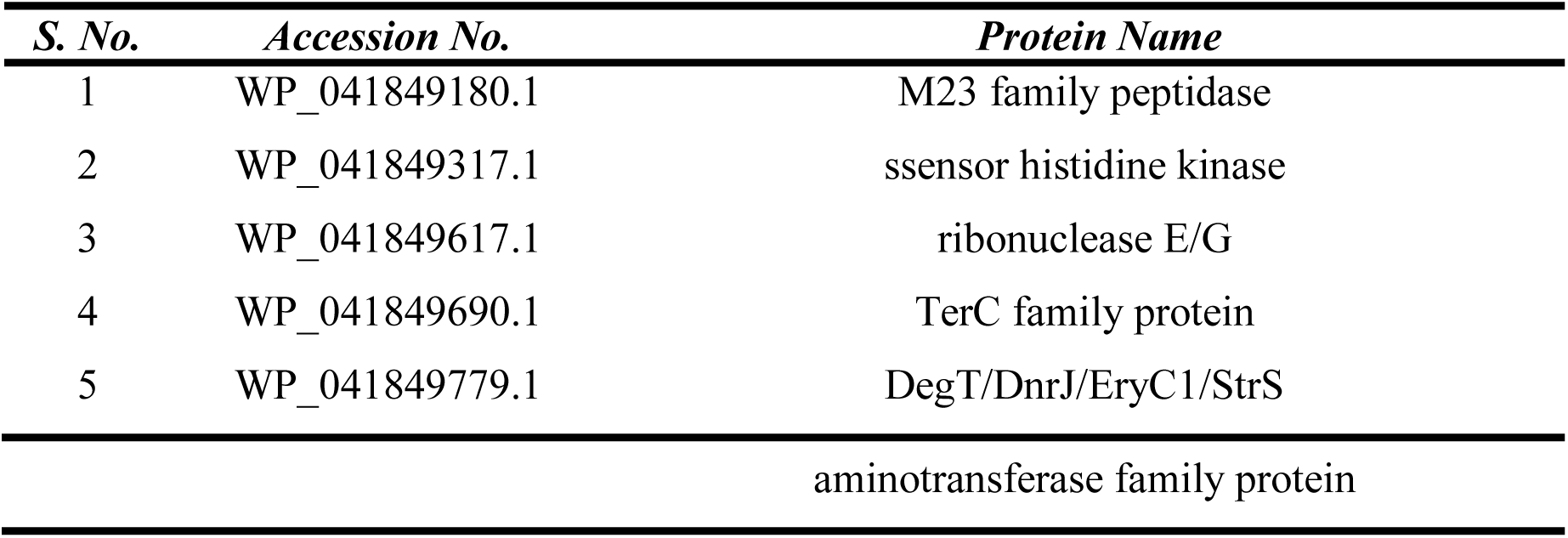
Proteins having no KO assignment.

### Subcellular Location Analysis of Proteins Involved in Pathogen-Specific Pathways

It was observed for many proteins which can span multiple localization sites and hence localization is one of the most important aspects of any therapeutic target [40]. Proteins listed in set4-a were passed through PSORTb v3.0.2, CELLO v.2.5 and ngLOC servers. Counting the best score for any predicted location for a protein, the final result were selected considering two conditions, 1) the indifferent location of protein predicted by all three servers, and we selected the locations of 6 proteins by this consideration; 2) the indifferent location predicted by any two servers and we selected the location of 1 protein by this consideration. 4 proteins were to be present in inner membrane while the rest 3 were found to be present in the cytoplasm. The results are provided in Supplementary Table 2. Excluding the cytoplasmic proteins, only inner membrane proteins were listed in set5 for further analysis and the accession numbers are provided in Table 5. More than 60% of therapeutic and drug targets are represented membrane proteins[41]. There are several advantages of targeting membrane proteins for treatment, for example, their function can easily be studied by computer-based tools and approaches rather than wet-lab processes. Computational tools can be used to predict the secondary structure since membrane proteins generally have a unique structure and show a propensity to form secondary structure[42]. In future, structure-based drug designing will be possible[43] by using ab initio modeling method, while no experimental information or 3D structure of high resolution is available[42,44].

**Table 5:** List of membrane proteins.

| *Accession No.* | *Protein* | *Location* |
| --- | --- | --- |
| WP_041849190.1 | histidine kinase | Inner Membrane |
| WP_041849727.1 | penicillin-binding protein 2 | Inner Membrane |
| WP_041849879.1 | flagellar biosynthesis protein FlhA | Inner Membrane |
| WP_052472869.1 | ABC transporter permease | Inner Membrane |

### Identification of Novel Drug Targets by Screening and druggability Analysis

Inner membrane proteins were screened through DrugBank Database to identify novelty of proteins which be used as drug targets. Two proteins showed similarity with the drug targets available at the database. They are provided in Table 6. Since our aim was to find novel drug targets we filtered these two proteins and listed the remaining two proteins in set5 which were found to be absent in the database. The broad spectrum therapeutic compounds medicated for a pathogenic strain or group of strains may transfer genes to other pathogenic strains as well as cause mutational changes. Due to overuse and misuse of antibiotics as well as the lack of designing new drugs by the pharmaceutical industry, the antibiotic resistance crisis has emerged[41,42]. To meet the challenge of this crisis, the selected novel drug targets in this can play a vital role in developing new therapeutic compounds.

**Table 6:**
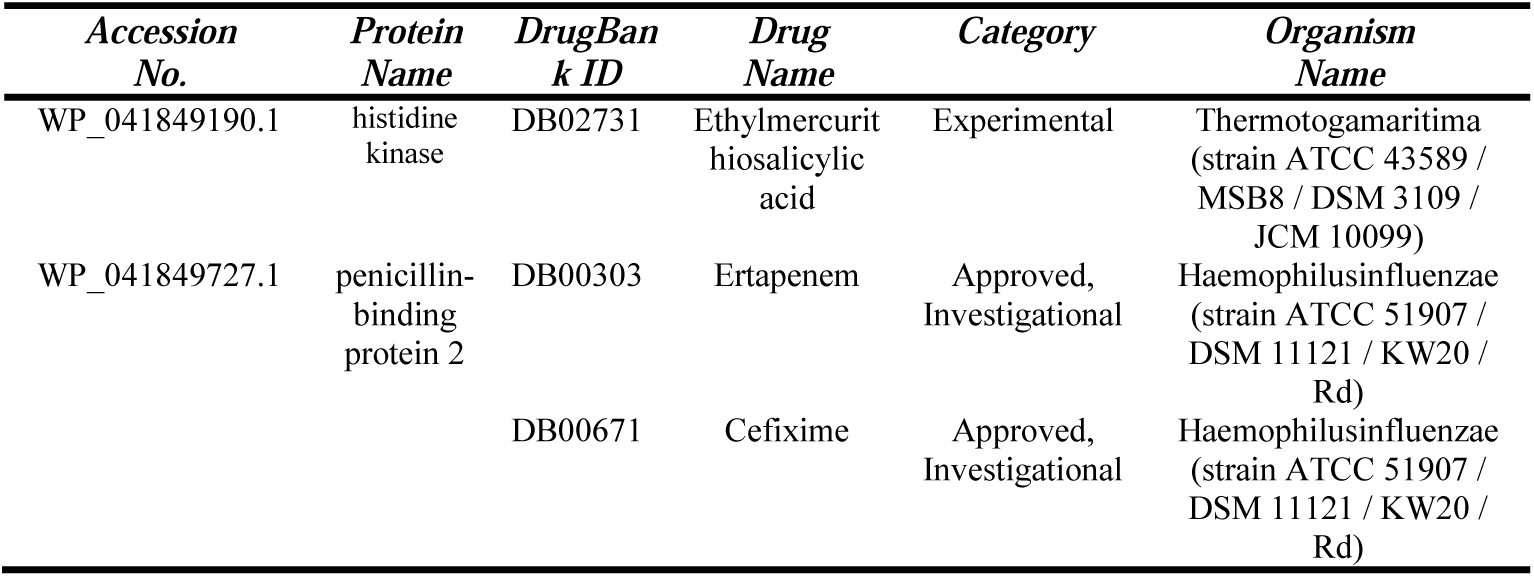
List of druggable proteins.

### ‘Anti-targets’ Analysis of Novel Drug Targets

Many drug candidates were withdrawn from markets due to showing carcinogenic effects, and that is why cross reactivity and carcinogenic checking is crucial for developing an effective drug molecule. [43-48]. Though host non-homologous proteins were removed from the the non-paralogous protein sequences of the pathogen, this analysis was performed to avoid any kind of cross-reactivity and toxic effects due to docking between the drugs administered for the pathogen and the host ‘anti-targets’. Set5 proteins were successfully screened through BLASTp with 210 reported ‘anti-targets’ present in human. No evidence of similarity was seen and hence all of set5 proteins were listed in set6 considering non-homologous to host ‘anti-targets’.

### Prediction of Antigenicity (Set8)

New genomics-based approaches or alternatively known as reverse vaccinology have been demonstrated to be an effective as well as powerful in discovering vaccine candidates [49-52]. Development of safe recombinant vaccines on the basis of the small number of antigenic protein sequences is emerging as the most attractive even cost-effective solution to combat with infectious disease [53-55]. Each novel drug target of *B. bacilliformis* subsp. ver097 in set7 was screened through VaxiJen v2.0 server. The server predicted ‘Flagellar biosynthesis protein FlhA (WP_041849879.1)’ and ‘ABC transporter permease (WP_052472869.1)’ as probable antigens of *B. bacilliformis* subsp. Ver097 and the result is provided in Table 7. Development of vaccine candidates against these membrane proteins can contribute to the control and eradication of infectious diseases of human which type of studies have already been reported [56,57].

**Table 7:** Novel drug target proteins showing antigenicity.

| *Accession No.* | *Protein Name* | *VaxiJenScore* |
| --- | --- | --- |
| WP_041849879.1 | flagellar biosynthesis protein FlhA | 0.6098 |
| WP_052472869.1 | ABC transporter permease | 0.4516 |

### Conservancy Analysis of Predicted Sequences with Other Strains

Protein-protein BLASTp revealed the conservancy pattern of ‘Flagellar biosynthesis protein FlhA (WP_041849879.1)’ and ‘ABC transporter permease (WP_052472869.1)’ from *B. bacilliformis* subsp. Ver097 with other strains of *B. bacilliformis*. ‘ABC transporter permease’ of *B. bacilliformis* subsp. Ver097 showed minimal conservancy with other *Bartonella bacilliformis* strains while ‘Flagellar biosynthesis protein FlhA’ of *B. bacilliformis* subsp. Ver097 showed above 99% conservancy with homologous protein of other classically used *Bartonella bacilliformis* strains (Table 8). Predicted protein always should be conserved in a wide range of bacterial strains for ensuring an effective drug target against a specific bacterial pathogen [58]. Here, ‘Flagellar biosynthesis protein FlhA’ displayed a higher conservancy pattern with homologous proteins of other *Bartonella bacilliformis* strains from different geographical locations. So, ‘Flagellar biosynthesis protein FlhA’ could be a potent broad spectrum drug target for *Bartonella bacilliformis.*

**Table 8:** Conservancy analysis Flagellar biosynthesis protein FlhA; (BLASTp results of WP_041849897.1 against other *Bartonellabacilliformis* strains)s

| *S. No.* | *Sequence id* | *Organism* | *Identity* |
| --- | --- | --- | --- |
| 1 | WP_005767817.1 | *Bartonellabacilliformis* KC583 | 99% |
| 2 | EKS43057.1 | *Bartonellabacilliformis* INS | 99% |
| 3 | EYS88603.1 | *Bartonellabacilliformis* San Pedro600-02 | 99% |
| 4 | EYS94419.1 | *Bartonellabacilliformis* Peru-18 | 99% |
| 5 | KEG15906.1 | *Bartonellabacilliformis* CUSCO5 | 99% |
| 6 | KEG19809.1 | *Bartonellabacilliformis* Peru38 | 99% |
| 7 | AMG86163.1 | *Bartonellabacilliformis* | 99% |
| 8 | KZM37516.1 | *Bartonellabacilliformis* | 99% |
| 9 | KZN21555.1 | *Bartonellabacilliformis* | 99% |
| 10 | KEG22213.1 | *Bartonellabacilliformis* Ver075 | 99% |
| 11 | KEG16163.1 | *Bartonellabacilliformis* Cond044 | 99% |
| 12 | EYS91027.1 | *Bartonellabacilliformis* str. Heidi Mejia | 99% |
| 13 | KEG18152.1 | *Bartonellabacilliformis* Hosp800-02 | 99% |
| 14 | KEG21780.1 | *Bartonellabacilliformis* VAB9028 | 99% |
| 15 | KEG23155.1 | *Bartonellabacilliformis* CAR600-02 | 99% |

## Conclusion

Subtractive genomic approach was successfully used throughout the study, and revealed that one protein of *Bartonella bacilliformis subsp. ver097*can be used as novel drug target and vaccine candidate. Inhibition of this inner membrane protein (Flagellar biosynthesis protein FlhA) will be helpful to combat against the infectious diseases caused by *Bartonella bacilliformis* pathogen since this protein is involved in pathogen-specific ‘flagellar assembly’ metabolic pathway. Furthermore chances of cross-reactivity between drugs and human proteins have been reduced since they showed no similarity against the human proteome and ‘anti-targets’. Hence the development of new vaccines and novel drugs or therapeutic compounds could be a promising footsteps to the eradication of diseases progession by *Bartonella bacilliformis.*

## Data Availability

Not Applicable

## Conflict of Interest

There is no conflict of interest regarding the publication of this paper

## Acknowledgment

We are highly thankful to Md. Ismail Hosen (Department of Biochemistry and Molecular Biology, University of Dhaka, Dhaka 1000, Bangladesh) for his cordial advice and suggestions.

## Funding source

There was no funding source for this project.

## Supplementary Materials

Supplementary file S1:’Anti-targets’ protein sequences (210) fetched from NCBI Protein database that can cause undesirable side effects during drug interactions.

## Supplementary Tables

Supplementary Table 1: List of unique metabolic pathways of the pathogen

Supplementary Table 2: Results of subcellular location of 7 non-homologous, essential proteins only involved in the unique metabolic pathways of the pathogen

