## Supplementary material for "Subtractive Genomic Analysis for Identification of Novel Drug Targets and Vaccine Candidates against *Bartonella bacilliformis* subsp. Ver097": Supplymentary Table 1

| S. No. |  | Pathway id | Unique Pathways |
| --- | --- | --- | --- |
| 1 |  | lpn00121 | Secondary bile acid biosynthesis |
| 2 |  | lpn00261 | Monobactam biosynthesis |
| 3 |  | lpn00281 | Geraniol degradation |
| 4 |  | lpn00300 | Lysine biosynthesis |
| 5 |  | lpn00332 | Carbapenem biosynthesis |
| 6 |  | lpn00361 | Chlorocyclohexane and chlorobenzene degradation |
| 7 |  | lpn00362 | Benzoate degradation |
| 8 |  | lpn00364 | Fluorobenzoate degradation |
| 9 |  | lpn00401 | Novobiocin biosynthesis |
| 10 |  | lpn00410 | beta-Alanine metabolism |
| 11 |  | lpn00460 | Cyanoamino acid metabolism |
| 12 |  | lpn00473 | D-Alanine metabolism |
| 13 |  | lpn00480 | Glutathione metabolism |
| 14 |  | lpn00521 | Streptomycin biosynthesis |
| 15 |  | lpn00523 | Polyketide sugar unit biosynthesis |
| 16 |  | lpn00525 | Acarbose and validamycin biosynthesis |
| 17 |  | lpn00540 | Lipopolysaccharide biosynthesis |
| 18 |  | lpn00623 | Toluene degradation |
| 19 |  | lpn00625 | Chloroalkane and chloroalkene degradation |
| 20 |  | lpn00626 | Naphthalene degradation |
| 21 |  | lpn00627 | Aminobenzoate degradation |
| 22 |  | lpn00643 | Styrene degradation |
| 23 |  | lpn00680 | Methane metabolism |
| 24 |  | lpn00903 | Limonene and pinene degradation |
| 25 |  | lpn00910 | Nitrogen metabolism |
| 26 |  | lpn00930 | Caprolactam degradation |
| 27 |  | lpn01110 | Biosynthesis of secondary metabolites |
| 28 |  | lpn01120 | Microbial metabolism in diverse environments |
| 29 |  | lpn01130 | Biosynthesis of antibiotics |
| 30 |  | lpn01220 | Degradation of aromatic compounds |
| 31 |  | lpn01501 | beta-Lactam resistance |
| 32 |  | lpn01502 | Vancomycin resistance |
| 33 |  | lpn01503 | Cationic antimicrobial peptide (CAMP) resistance |
| 34 |  | lpn02020 | Two-component system |
| 35 |  | lpn02024 | Quorum sensing |
| 36 |  | lpn02030 | Bacterial chemotaxis |
| 37 |  | lpn02040 | Flagellar assembly |
| 38 |  | lpn03060 | Protein export |
| 39 |  | lpn03070 | Bacterial secretion system |
| 40 |  | lpn03410 | Base excision repair |
| 41 |  | lpn00550 | Peptidoglycan biosynthesis |
