## Supplementary material for "Subtractive Genomic Analysis for Identification of Novel Drug Targets and Vaccine Candidates against *Bartonella bacilliformis* subsp. Ver097": Supplymentary Table 2

| S. no. | Accession No. | PSORTb v3.0.2 server | CELLO v2.5 server | ngLOC server | PSLpred server | Fianl result |
| --- | --- | --- | --- | --- | --- | --- |
| 1 | tr\|Q5ZXH3 | Cytoplasmic | Extracellular | Inner Membrane | Cytoplasmic | Cytoplasmic |
| 2 | tr\|Q5ZWF1 | Cytoplasmic | Cytoplasmic | Cytoplasmic |  | Cytoplasmic |
| 3 | tr\|Q5ZTC6 | Cytoplasmic  Membrane | Inner Membrane \| | Cytoplasmic |  | Inner Membranene \| |
| 4 | tr\|Q5ZSM2 | Unknown | Cytoplasmic | Cytoplasmic |  | Cytoplasmic |
| 5 | tr\|Q5ZXL1 | Cytoplasmic  Membrane | Inner Membrane \| | Inner Membrane \| |  | Inner Membranene |
| 6 | tr\|Q5ZW84 | Cytoplasmic  Membrane | Inner Membrane \| | Inner Membrane \| |  | Inner Membranene |
| 7 | sp\|Q5ZZK8 | Cytoplasmic | Cytoplasmic | Cytoplasmic |  |  |
| 8 | tr\|Q5ZT47 | Cytoplasmic  Membrane | Inner Membrane \| | Inner Membrane \| |  | Inner Membranene |
| 9 | tr\|Q5ZY95 | Cytoplasmic | Cytoplasmic | Cytoplasmic |  | Cytoplasmic |
| 10 | sp\|Q5ZVH7 | Multiple location | Cytoplasmic | Inner Membrane | Cytoplasmic | Cytoplasmic |
| 11 | tr\|Q5ZYM3 | Cytoplasmic  Membrane | Inner Membrane \| | Inner Membrane \| |  | Inner Membranene |
| 12 | tr\|Q3V868 | Cytoplasmic  Membrane | Inner Membrane \| | Inner Membrane \| |  | Inner Membrane |
| 13 | tr\|Q5ZU07 | Cytoplasmic  Membrane | Inner Membrane \| | Inner Membrane \| |  | Inner Membrane \| |
| 14 | tr\|Q5ZU08 | Cytoplasmic  Membrane | Inner Membrane \| | Inner Membrane |  | Inner Membrane \| |
| 15 | tr\|Q5ZVW9 | Cytoplasmic | Cytoplasmic | Cytoplasmic |  | Cytoplasmic |
| 16 | tr\|Q5ZRK1 | Cytoplasmic | Cytoplasmic | Cytoplasmic |  | Cytoplasmic |
| 17 | tr\|Q5ZX17 | Cytoplasmic  Membrane | Inner Membrane \| | Cytoplasmic |  | Inner Membrane \| |
| 18 | tr\|Q5ZVR5 | Cytoplasmic  Membrane | Outer Membrane | Inner Membrane \| |  | Inner Membrane \| |
| 19 | tr\|Q5ZX16 | Cytoplasmic | Cytoplasmic | Cytoplasmic |  | Cytoplasmic |
