## Supplementary material for "Subtractive Genomic Analysis for Identification of Novel Drug Targets and Vaccine Candidates against *Bartonella bacilliformis* subsp. Ver097": Supplymentary-file-S1

**Supplementary file S1:‘**Anti-targets’ protein sequences (210) fetched from NCBI Protein database that can cause undesirable side effects during drug interactions.

>gi|13236497|ref|NP_076917.1| 5-hydroxytryptamine receptor 5A [Homo sapiens]

>gi|1168243|sp|P25100.2|ADA1D_HUMAN Alpha-1D adrenergic receptor (Alpha 1D-adrenoreceptor) (Alpha 1D-adrenoceptor) (Alpha-1A adrenergic receptor) (Alpha-adrenergic receptor 1a)

>gi|1168246|sp|P35348.2|ADA1A_HUMAN Alpha-1A adrenergic receptor (Alpha 1A-adrenoreceptor) (Alpha 1A-adrenoceptor) (Alpha-1C adrenergic receptor) (Alpha-adrenergic receptor 1c)

>gi|7690135|gb|AAB31163.2| alpha adrenergic receptor subtype alpha 1a [Homo sapiens]

>gi|547222|gb|AAB31165.1| alpha adrenergic receptor subtype alpha 1c [human, heart, Peptide, 466 aa]

>gi|547221|gb|AAB31164.1| alpha adrenergic receptor subtype alpha 1b [human, heart, Peptide, 516 aa]

>gi|2978556|gb|AAC06138.1| alpha 1A adrenergic receptor isoform 4 [Homo sapiens]

>gi|52000961|sp|P63092.1|GNAS2_HUMAN Guanine nucleotide-binding protein G(s) subunit alpha isoforms short (Adenylate cyclase-stimulating G alpha protein)

>gi|148539879|ref|NP_005151.2| beta adrenergic receptor kinase 2 [Homo sapiens]

>gi|15004694|gb|AAK77197.1|AF395806_1 adrenergic receptor alpha-1a [Homo sapiens]

>gi|111118990|ref|NP_150647.2| alpha-1A-adrenergic receptor isoform 4 [Homo sapiens]

>gi|111118988|ref|NP_150645.2| alpha-1A-adrenergic receptor isoform 3 [Homo sapiens]

>gi|111118986|ref|NP_150646.3| alpha-1A-adrenergic receptor isoform 2 [Homo sapiens]

>gi|4501959|ref|NP_000670.1| alpha-1B-adrenergic receptor [Homo sapiens]

>gi|177807|gb|AAA35496.1| alpha-1A-adrenergic receptor

>gi|117938756|ref|NP_001070950.1| constitutive androstane receptor isoform 1 [Homo sapiens]

>gi|117938754|ref|NP_001070949.1| constitutive androstane receptor isoform 4 [Homo sapiens]

>gi|117938752|ref|NP_001070948.1| constitutive androstane receptor isoform 2 [Homo sapiens]

>gi|117938750|ref|NP_001070947.1| constitutive androstane receptor isoform 10 [Homo sapiens]

>gi|117938748|ref|NP_001070945.1| constitutive androstane receptor isoform 14 [Homo sapiens]

>gi|117938746|ref|NP_001070946.1| constitutive androstane receptor isoform 7 [Homo sapiens]

>gi|117938744|ref|NP_001070944.1| constitutive androstane receptor isoform 13 [Homo sapiens]

>gi|117938742|ref|NP_001070943.1| constitutive androstane receptor isoform 15 [Homo sapiens]

>gi|117938740|ref|NP_001070942.1| constitutive androstane receptor isoform 8 [Homo sapiens]

>gi|117938738|ref|NP_001070941.1| constitutive androstane receptor isoform 12 [Homo sapiens]

>gi|117938736|ref|NP_001070940.1| constitutive androstane receptor isoform 9 [Homo sapiens]

>gi|117938734|ref|NP_001070939.1| constitutive androstane receptor isoform 5 [Homo sapiens]

>gi|117938732|ref|NP_001070938.1| constitutive androstane receptor isoform 11 [Homo sapiens]

>gi|117938729|ref|NP_001070937.1| constitutive androstane receptor isoform 6 [Homo sapiens]

>gi|4826661|ref|NP_005113.1| constitutive androstane receptor isoform 3 [Homo sapiens]

>gi|48869156|gb|AAT47173.1| constitutive androstane receptor SV15 [Homo sapiens]

>gi|48869154|gb|AAT47172.1| constitutive androstane receptor SV14 [Homo sapiens]

>gi|48869152|gb|AAT47171.1| constitutive androstane receptor SV13 [Homo sapiens]

>gi|48869150|gb|AAT47170.1| constitutive androstane receptor SV12 [Homo sapiens]

>gi|48869148|gb|AAT47169.1| constitutive androstane receptor SV11 [Homo sapiens]

>gi|48869142|gb|AAT47166.1| constitutive androstane receptor SV8 [Homo sapiens]

>gi|48869140|gb|AAT47165.1| constitutive androstane receptor SV7 [Homo sapiens]

>gi|48869128|gb|AAT47159.1| constitutive androstane receptor SV1 [Homo sapiens]

>gi|48869168|gb|AAT47179.1| constitutive androstane receptor SV21 [Homo sapiens]

>gi|48869166|gb|AAT47178.1| constitutive androstane receptor SV20 [Homo sapiens]

>gi|48869162|gb|AAT47176.1| constitutive androstane receptor SV18 [Homo sapiens]

>gi|48869146|gb|AAT47168.1| constitutive androstane receptor SV10 [Homo sapiens]

>gi|48869158|gb|AAT47174.1| constitutive androstane receptor SV16 [Homo sapiens]

>gi|48869130|gb|AAT47160.1| constitutive androstane receptor SV2 [Homo sapiens]

>gi|32307128|ref|NP_054790.2| nuclear receptor coactivator 6 [Homo sapiens]

>gi|52630419|ref|NP_001005291.1| sterol regulatory element binding transcription factor 1 isoform a [Homo sapiens]

>gi|22547195|ref|NP_004167.3| sterol regulatory element binding transcription factor 1 isoform b [Homo sapiens]

>gi|23110952|ref|NP_683865.1| cathepsin E isoform b preproprotein [Homo sapiens]

>gi|4503145|ref|NP_001901.1| cathepsin E isoform a preproprotein [Homo sapiens]

>gi|38327531|ref|NP_938151.1| insulin induced gene 1 isoform 3 [Homo sapiens]

>gi|38327529|ref|NP_938150.1| insulin induced gene 1 isoform 2 [Homo sapiens]

>gi|28882053|ref|NP_005533.2| insulin induced gene 1 isoform 1 [Homo sapiens]

>gi|193083117|ref|NP_001122387.1| cytochrome P450 family 24 subfamily A polypeptide 1 isoform 2 precursor [Homo sapiens]

>gi|55770850|ref|NP_000773.2| cytochrome P450 family 24 subfamily A polypeptide 1 isoform 1 precursor [Homo sapiens]

>gi|46249376|ref|NP_996759.1| protein phosphatase 1, catalytic subunit, beta isoform 1 [Homo sapiens]

>gi|29540545|ref|NP_003158.2| sulfotransferase family, cytosolic, 2A, dehydroepiandrosterone-preferring, member 1 [Homo sapiens]

>gi|117205|sp|P20813.1|CP2B6_HUMAN Cytochrome P450 2B6 (CYPIIB6) (P450 IIB1)

>gi|5921480|emb|CAB56463.1| dopamine receptor D2 [Homo sapiens]

>gi|4467834|emb|CAB37869.1| dopamine receptor D2 [Homo sapiens]

>gi|1345939|sp|P21917.2|DRD4_HUMAN D(4) dopamine receptor (Dopamine D4 receptor) (D(2C) dopamine receptor)

>gi|17986270|ref|NP_057658.2| dopamine receptor D2 isoform short [Homo sapiens]

>gi|405310|gb|AAB26819.1| D2 dopamine receptor [Homo sapiens]

>gi|553269|gb|AAA52328.1| dopamine D2 receptor

>gi|3820492|gb|AAC78779.1| dopamine D2 receptor [Homo sapiens]

>gi|62089250|dbj|BAD93069.1| dopamine receptor D2 isoform long variant [Homo sapiens]

>gi|119587629|gb|EAW67225.1| dopamine receptor D2, isoform CRA_e [Homo sapiens]

>gi|119587627|gb|EAW67223.1| dopamine receptor D2, isoform CRA_d [Homo sapiens]

>gi|119587623|gb|EAW67219.1| dopamine receptor D2, isoform CRA_a [Homo sapiens]

>gi|32483397|ref|NP_000788.2| dopamine receptor D4 [Homo sapiens]

>gi|4503383|ref|NP_000785.1| dopamine receptor D1 [Homo sapiens]

>gi|89191861|ref|NP_000787.2| dopamine receptor D3 isoform a [Homo sapiens]

>gi|89191863|ref|NP_387512.3| dopamine receptor D3 isoform e [Homo sapiens]

>gi|7381416|gb|AAF61479.1|AF176812_1 dopamine receptor D2longer [Homo sapiens]

>gi|5921992|ref|NP_000666.2| adenosine A2a receptor [Homo sapiens]

>gi|12548788|ref|NP_068712.1| gamma-aminobutyric acid (GABA) A receptor, beta 3 isoform 2 precursor [Homo sapiens]

>gi|4503867|ref|NP_000805.1| gamma-aminobutyric acid (GABA) A receptor, beta 3 isoform 1 precursor [Homo sapiens]

>gi|157389018|ref|NP_000802.2| gamma-aminobutyric acid A receptor, alpha 6 precursor [Homo sapiens]

>gi|150378528|ref|NP_001092881.1| G protein-coupled receptor associated sorting protein 1 [Homo sapiens]

>gi|4507685|ref|NP_003295.1| transient receptor potential cation channel, subfamily C, member 1 [Homo sapiens]

>gi|20373106|dbj|BAB91222.1| muscarinic acetylcholine receptor M5 [Homo sapiens]

>gi|30425444|ref|NP_848605.1| ankyrin repeat and kinase domain containing 1 [Homo sapiens]

>gi|66933005|ref|NP_001019820.1| calnexin precursor [Homo sapiens]

>gi|4504041|ref|NP_002061.1| guanine nucleotide binding protein (G protein), alpha inhibiting activity polypeptide 2 [Homo sapiens]

>gi|50541890|gb|AAT78421.1| Galphai2 protein [Homo sapiens]

>gi|4885563|ref|NP_005391.1| protein kinase C, epsilon [Homo sapiens]

>gi|96974985|ref|NP_060191.3| coiled-coil and C2 domain containing 1A [Homo sapiens]

>gi|86990435|ref|NP_001034556.1| G protein signalling regulator 19 [Homo sapiens]

>gi|34452715|ref|NP_003707.2| Ca2+-dependent secretion activator isoform 1 [Homo sapiens]

>gi|34452713|ref|NP_899631.1| Ca2+-dependent secretion activator isoform 2 [Homo sapiens]

>gi|34452711|ref|NP_899630.1| Ca2+-dependent secretion activator isoform 3 [Homo sapiens]

>gi|4758020|ref|NP_000605.1| ciliary neurotrophic factor [Homo sapiens]

>gi|85540964|sp|Q86UW7.2|CAPS2_HUMAN Calcium-dependent secretion activator 2 (Calcium-dependent activator protein for secretion 2) (CAPS-2)

>gi|192447423|ref|NP_001122298.1| frequenin homolog isoform 2 [Homo sapiens]

>gi|17738308|ref|NP_055101.2| frequenin homolog isoform 1 [Homo sapiens]

>gi|140972063|ref|NP_115984.3| protein phosphatase 1, regulatory subunit 9B [Homo sapiens]

>gi|148839284|ref|NP_001009571.2| Ca2+-dependent activator protein for secretion 2 isoform b [Homo sapiens]

>gi|563756|gb|AAA88023.1| unknown protein

>gi|37622910|ref|NP_000729.2| cholinergic receptor, muscarinic 1 [Homo sapiens]

>gi|3292965|emb|CAA07081.1| m1 muscarinic acetylcholine receptor protein [Homo sapiens]

>gi|14573537|gb|AAK68112.1| m1 muscarinic cholinergic receptor [Homo sapiens]

>gi|14285389|sp|O43526.2|KCNQ2_HUMAN Potassium voltage-gated channel subfamily KQT member 2 (Voltage-gated potassium channel subunit Kv7.2) (Neuroblastoma-specific potassium channel subunit alpha KvLQT2) (KQT-like 2)

>gi|5921785|sp|O43525.2|KCNQ3_HUMAN Potassium voltage-gated channel subfamily KQT member 3 (Voltage-gated potassium channel subunit Kv7.3) (Potassium channel subunit alpha KvLQT3) (KQT-like 3)

>gi|6166005|sp|P51787.3|KCNQ1_HUMAN Potassium voltage-gated channel subfamily KQT member 1 (Voltage-gated potassium channel subunit Kv7.1) (IKs producing slow voltage-gated potassium channel subunit alpha KvLQT1) (KQT-like 1)

>gi|6166006|sp|P56696.1|KCNQ4_HUMAN Potassium voltage-gated channel subfamily KQT member 4 (Voltage-gated potassium channel subunit Kv7.4) (Potassium channel subunit alpha KvLQT4) (KQT-like 4)

>gi|41327726|ref|NP_000738.2| nicotinic acetylcholine receptor beta 1 subunit precursor [Homo sapiens]

>gi|40254462|ref|NP_002063.2| guanine nucleotide binding protein (G protein), q polypeptide [Homo sapiens]

>gi|26638655|ref|NP_751895.1| potassium voltage-gated channel KQT-like protein 4 isoform b [Homo sapiens]

>gi|27886590|ref|NP_775268.1| PTK2B protein tyrosine kinase 2 beta isoform a [Homo sapiens]

>gi|27886588|ref|NP_775267.1| PTK2B protein tyrosine kinase 2 beta isoform b [Homo sapiens]

>gi|51093859|ref|NP_001003406.1| calcium channel, voltage-dependent, T type, alpha 1I subunit isoform b [Homo sapiens]

>gi|21361077|ref|NP_066919.2| calcium channel, voltage-dependent, T type, alpha 1I subunit isoform a [Homo sapiens]

>gi|115511049|ref|NP_002058.2| guanine nucleotide binding protein (G protein), alpha 11 (Gq class) [Homo sapiens]

>gi|6093860|sp|O75469.1|NR1I2_HUMAN Nuclear receptor subfamily 1 group I member 2 (Orphan nuclear receptor PXR) (Pregnane X receptor) (Orphan nuclear receptor PAR1) (Steroid and xenobiotic receptor) (SXR)

>gi|14702164|ref|NP_148934.1| pregnane X receptor isoform 3 [Homo sapiens]

>gi|11863132|ref|NP_071285.1| pregnane X receptor isoform 2 [Homo sapiens]

>gi|126031433|pdb|2O9I|B Chain B, Crystal Structure Of The Human Pregnane X Receptor Lbd In Complex With An Src-1 Coactivator Peptide And T0901317

>gi|14719569|pdb|1ILH|A Chain A, Crystal Structure Of Human Pregnane X Receptor Ligand Binding Domain Bound To Sr12813

>gi|22538457|ref|NP_671756.1| nuclear receptor coactivator 1 isoform 2 [Homo sapiens]

>gi|22538459|ref|NP_671766.1| nuclear receptor coactivator 1 isoform 3 [Homo sapiens]

>gi|22538455|ref|NP_003734.3| nuclear receptor coactivator 1 isoform 1 [Homo sapiens]

>gi|13752550|gb|AAK38720.1|AF364606_1 orphan nuclear receptor PXR.1 [Homo sapiens]

>gi|71725341|ref|NP_001025174.1| hepatocyte nuclear factor 4 alpha isoform e [Homo sapiens]

>gi|71725339|ref|NP_787110.2| hepatocyte nuclear factor 4 alpha isoform d [Homo sapiens]

>gi|71725336|ref|NP_001025175.1| hepatocyte nuclear factor 4 alpha isoform f [Homo sapiens]

>gi|31077209|ref|NP_849181.1| hepatocyte nuclear factor 4 alpha isoform c [Homo sapiens]

>gi|31077207|ref|NP_849180.1| hepatocyte nuclear factor 4 alpha isoform a [Homo sapiens]

>gi|31077205|ref|NP_000448.3| hepatocyte nuclear factor 4 alpha isoform b [Homo sapiens]

>gi|62737968|pdb|1SKX|A Chain A, Structural Disorder In The Complex Of Human Pxr And The Macrolide Antibiotic Rifampicin

>gi|61679486|pdb|1XV9|D Chain D, Crystal Structure Of CarRXR HETERODIMER BOUND WITH SRC1 Peptide, Fatty Acid, And 5b-Pregnane-3,20-Dione.

>gi|61679485|pdb|1XV9|C Chain C, Crystal Structure Of CarRXR HETERODIMER BOUND WITH SRC1 Peptide, Fatty Acid, And 5b-Pregnane-3,20-Dione.

>gi|10863829|gb|AAG23345.1| PAR2 [Homo sapiens]

>gi|307180|gb|AAA59575.1| P-glycoprotein [Homo sapiens]

>gi|386862|gb|AAA59576.1| P glycoprotein

>gi|34523|emb|CAA41558.1| P-glycoprotein [Homo sapiens]

>gi|2394178|gb|AAB70218.1| P-glycoprotein [Homo sapiens]

>gi|2353264|gb|AAB69423.1| P-glycoprotein [Homo sapiens]

>gi|33307712|gb|AAQ03033.1|AF399931_1 P-glycoprotein [Homo sapiens]

>gi|37543520|gb|AAM09027.1| P-glycoprotein [Homo sapiens]

>gi|25990364|gb|AAN76500.1|AF319622_1 P-glycoprotein [Homo sapiens]

>gi|307181|gb|AAA36207.1| P-glycoprotein

>gi|41058415|gb|AAR99172.1| P-glycoprotein [Homo sapiens]

>gi|2506118|sp|P08183.2|MDR1_HUMAN Multidrug resistance protein 1 (ATP-binding cassette sub-family B member 1) (P-glycoprotein 1) (CD243 antigen)

>gi|126302568|sp|P21439.2|MDR3_HUMAN Multidrug resistance protein 3 (ATP-binding cassette sub-family B member 4) (P-glycoprotein 3)

>gi|1096242|prf||2111304A P glycoprotein

>gi|1006663|emb|CAA84542.1| MDR3 P-glycoprotein [Homo sapiens]

>gi|34525|emb|CAA29547.1| P-glycoprotein (431 AA) [Homo sapiens]

>gi|6681436|dbj|BAA88711.1| sister p-glycoprotein [Homo sapiens]

>gi|116242940|sp|Q2M3G0.2|ABCB5_HUMAN ATP-binding cassette sub-family B member 5 (P-glycoprotein ABCB5) (ABCB5 P-gp)

>gi|40795903|gb|AAR91622.1| P-glycoprotein 1 [Homo sapiens]

>gi|9961252|ref|NP_061338.1| ATP-binding cassette, subfamily B, member 4 isoform C [Homo sapiens]

>gi|21536378|ref|NP_003733.2| ATP-binding cassette, sub-family B (MDR/TAP), member 11 [Homo sapiens]

>gi|4506201|ref|NP_002788.1| proteasome beta 5 subunit [Homo sapiens]

>gi|47116933|sp|Q9NS86.1|LANC2_HUMAN LanC-like protein 2 (Testis-specific adriamycin sensitivity protein)

>gi|19913412|ref|NP_005106.2| major vault protein [Homo sapiens]

>gi|25777707|ref|NP_740753.1| zinc ribbon domain containing 1 [Homo sapiens]

>gi|66529005|ref|NP_005679.2| ATP-binding cassette, sub-family C, member 5 isoform 1 [Homo sapiens]

>gi|66529093|ref|NP_001018881.1| ATP-binding cassette, sub-family C, member 5 isoform 2 [Homo sapiens]

>gi|40287613|gb|AAR83914.1| unknown [Homo sapiens]

>gi|11545827|ref|NP_071395.1| p53-regulated apoptosis-inducing protein 1 [Homo sapiens]

>gi|121746|sp|P09211.2|GSTP1_HUMAN Glutathione S-transferase P (GST class-pi) (GSTP1-1)

>gi|56549095|ref|NP_001008406.1| B-cell receptor-associated protein BAP29 isoform c [Homo sapiens]

>gi|56549093|ref|NP_001008405.1| B-cell receptor-associated protein BAP29 isoform a [Homo sapiens]

>gi|56549091|ref|NP_061332.2| B-cell receptor-associated protein BAP29 isoform b [Homo sapiens]

>gi|4507811|ref|NP_003349.1| ceramide glucosyltransferase [Homo sapiens]

>gi|19913401|ref|NP_005066.2| organic anion transporting polypeptide A isoform b [Homo sapiens]

>gi|6005747|ref|NP_009143.1| ring finger protein 2 [Homo sapiens]

>gi|5729810|ref|NP_006570.1| emopamil binding protein (sterol isomerase) [Homo sapiens]

>gi|33946329|ref|NP_005393.2| ras related v-ral simian leukemia viral oncogene homolog A [Homo sapiens]

>gi|8923198|ref|NP_060182.1| pyroglutamyl-peptidase I [Homo sapiens]

>gi|19913403|ref|NP_602307.1| organic anion transporting polypeptide A isoform a [Homo sapiens]

>gi|19923262|ref|NP_004153.2| RAB5A, member RAS oncogene family [Homo sapiens]

>gi|2664295|emb|CAA84543.1| multidrug resistance protein 3 [Homo sapiens]

>gi|35553|emb|CAA41416.1| 70kDa peroxisomal membrane protein [Homo sapiens]

>gi|19584229|emb|CAD28599.1| unnamed protein product [Homo sapiens]

>gi|9955963|ref|NP_005680.1| ATP-binding cassette, sub-family B, member 6 [Homo sapiens]

>gi|11245446|gb|AAG33618.1| ATP-binding cassette half-transporter [Homo sapiens]

>gi|11245444|gb|AAG33617.1|AF308472_1 ATP-binding cassette half-transporter [Homo sapiens]

>gi|4156239|dbj|BAA37096.1| HERG [Homo sapiens]

>gi|37499017|gb|AAQ91594.1| potassium channel HERG [Homo sapiens]

>gi|37499029|gb|AAQ91600.1| potassium channel HERG [Homo sapiens]

>gi|37499027|gb|AAQ91599.1| potassium channel HERG [Homo sapiens]

>gi|26051273|ref|NP_742054.1| voltage-gated potassium channel, subfamily H, member 2 isoform c [Homo sapiens]

>gi|26051271|ref|NP_742053.1| voltage-gated potassium channel, subfamily H, member 2 isoform b [Homo sapiens]

>gi|118341648|gb|AAI27674.1| KCNH2 protein [Homo sapiens]

>gi|45945867|gb|AAH01914.2| KCNH2 protein [Homo sapiens]

>gi|34783890|gb|AAH04311.2| KCNH2 protein [Homo sapiens]

>gi|11933152|dbj|BAB19682.1| HERG-USO [Homo sapiens]

>gi|3549259|gb|AAC69709.1| HERG-USO [Homo sapiens]

>gi|157880355|pdb|1UJL|A Chain A, Solution Structure Of The Herg K+ Channel S5-P Extracellular Linker

>gi|119578199|gb|EAW57795.1| modifier of the HERG potassium channel [Homo sapiens]

>gi|52630440|ref|NP_036313.3| FK506-binding protein 8 [Homo sapiens]

>gi|31563383|ref|NP_733827.2| serum/glucocorticoid regulated kinase 3 isoform 2 [Homo sapiens]

>gi|75813626|ref|NP_001028750.1| serum/glucocorticoid regulated kinase 3 isoform 1 [Homo sapiens]

>gi|6685661|sp|Q9Y6J6.1|KCNE2_HUMAN Potassium voltage-gated channel subfamily E member 2 (Minimum potassium ion channel-related peptide 1) (Potassium channel subunit beta MiRP1) (MinK-related peptide 1)

>gi|116416|sp|P15382.1|KCNE1_HUMAN Potassium voltage-gated channel subfamily E member 1 (IKs producing slow voltage-gated potassium channel subunit beta Mink) (Minimal potassium channel) (Delayed rectifier potassium channel subunit IsK)

>gi|19923352|ref|NP_006158.2| NK3 homeobox 1 [Homo sapiens]

>gi|27436978|ref|NP_751951.1| potassium voltage-gated channel, Isk-related family, member 2 [Homo sapiens]

>gi|5803225|ref|NP_006752.1| tyrosine 3/tryptophan 5 -monooxygenase activation protein, epsilon polypeptide [Homo sapiens]

>gi|103488986|gb|ABF71886.1| voltage-gated potassium channel KV11.1 transcript variant 1 [Homo sapiens]

>gi|38635441|emb|CAE82156.1| potassium voltage-gated channel, subfamily H (eag-related), member 2 [Homo sapiens]

>gi|26006811|sp|Q9NS40.1|KCNH7_HUMAN Potassium voltage-gated channel subfamily H member 7 (Voltage-gated potassium channel subunit Kv11.3) (Ether-a-go-go-related gene potassium channel 3) (HERG-3) (Ether-a-go-go-related protein 3) (Eag-related protein 3)

>gi|26006810|sp|Q9H252.1|KCNH6_HUMAN Potassium voltage-gated channel subfamily H member 6 (Voltage-gated potassium channel subunit Kv11.2) (Ether-a-go-go-related gene potassium channel 2) (Ether-a-go-go-related protein 2) (Eag-related protein 2)

>gi|4033376|sp|Q14524.1|SCN5A_HUMAN Sodium channel protein type 5 subunit alpha (Sodium channel protein type V subunit alpha) (Voltage-gated sodium channel subunit alpha Nav1.5) (Sodium channel protein cardiac muscle subunit alpha) (HH1)

>gi|154448884|ref|NP_542402.2| potassium voltage-gated channel, Isk-related family, member 4 [Homo sapiens]

>gi|27886665|ref|NP_775185.1| potassium voltage-gated channel, subfamily H, member 7 isoform 2 [Homo sapiens]

>gi|4885443|ref|NP_005463.1| potassium voltage-gated channel, Isk-related family, member 3 [Homo sapiens]

>gi|27886651|ref|NP_775115.1| potassium voltage-gated channel, subfamily H, member 6 isoform 2 [Homo sapiens]

>gi|6729769|pdb|1BYW|A Chain A, Structure Of The N-Terminal Domain Of The Human-Erg Potassium Channel

>gi|71739466|gb|AAZ40507.1| potassium channel HERG1 [Homo sapiens]

>gi|3452413|emb|CAA09232.1| ether-a-go-go-related protein [Homo sapiens]
